# BitterSweet: Building machine learning models for predicting the bitter and sweet taste of small molecules

**DOI:** 10.1101/426692

**Authors:** Rudraksh Tuwani, Somin Wadhwa, Ganesh Bagler

## Abstract

The dichotomy of sweet and bitter tastes is a salient evolutionary feature of human gustatory system with an innate attraction to sweet taste and aversion to bitterness. A better understanding of molecular correlates of bitter-sweet taste gradient is crucial for identification of natural as well as synthetic compounds of desirable taste on this axis. While previous studies have advanced our understanding of the molecular basis of bitter-sweet taste and contributed models for their identification, there is ample scope to enhance these models by meticulous compilation of bitter-sweet molecules and utilization of a wide spectrum of molecular descriptors. Towards these goals, based on structured data compilation our study provides an integrative framework with state-of-the-art machine learning models for bitter-sweet taste prediction (BitterSweet). We compare different sets of molecular descriptors for their predictive performance and further identify important features as well as feature blocks. The utility of BitterSweet models is demonstrated by taste prediction on large specialized chemical sets such as FlavorDB, FooDB, SuperSweet, Super Natural II, DSSTox, and DrugBank. To facilitate future research in this direction, we make all datasets and BitterSweet models publicly available, and also present an end-to-end software for bitter-sweet taste prediction based on freely available chemical descriptors.

## 1. INTRODUCTION

Perception of taste is a complex sensation evolved in humans primarily to respond to naturally occurring food-derived chemicals^1^. Among all the taste perceptions, the dichotomy of sweet and bitter tastes is a salient evolutionary feature of the human gustatory system. The sweet taste is innately attractive, whereas bitterness evokes an aversive response. Receptors T1R2 and T1R3 belonging to the family of G-protein coupled receptors are known to be involved in the sensation of sweetness^2^. Interestingly, the bitterness sensation involves 25 hTAS2R receptors from the same repertoire of signaling proteins^3^. The sensation of bitter-sweet taste stems from complex interactions of a compound with these receptors. In addition to the oral cavity, taste receptors are present in other parts of the body such as the gut, respiratory system, and pancreas^1^. Beyond their primary role in taste perception (in the oral cavity), receptors in such locations are reported to be linked to mechanisms of diabetes and obesity by virtue of their involvement in nutrient perception, glucose level maintenance, appetite regulation and secretion of hormones^4-7^. Identification of compounds with a desirable gradient of bitter-sweet taste has immediate applications for developing low-calorie sweeteners and bitter masking molecules^8,9^. Thus, a better understanding of molecular correlates that are responsible for the bitter-sweet taste is of key value towards the identification of natural as well as synthetic compounds of desirable taste on this axis.

The mechanisms of gustatory sensation hinges on the structure of the receptor and that of the compounds. The perception of taste is highly sensitive to variations in compound structure, with subtle changes leading to a radical shift in the taste^10^. Moreover, completely resolved structures of sensory receptors are not available, further adding to the challenges of taste prediction. While ligand-based methods have found some success^11^, they have been restricted largely to specialized chemical families. With the availability of taste information of compounds and their molecular descriptors, data-driven approaches for building computational models towards prediction of taste are of immense value.

Previous studies have largely focussed on the problem of either bitter/non-bitter or sweet/non-sweet taste prediction, ignoring the dichotomy of bitter-sweet taste. In one of the pioneering studies for bitter-taste prediction, Rodgers et al.^12^ used a proprietary dataset of bitter molecules and randomly selected molecules (expected to be non-bitter), to develop a Naïve Bayes classifier utilizing circular fingerprints as molecular descriptors. Similarly, BitterX^13^ used random molecules to form their negative set while utilizing (publicly available) BitterDB^14^ compounds to form their positive set. As opposed to the use of 2D fingerprints, BitterX used physicochemical features of molecules (from Handbook of Molecular Descriptors^15^) towards the training of Support Vector Machine (SVM) classifier. Rojas et al.^16^ instead tackled the problem of sweet-taste prediction using experimentally verified data from the literature, 2D molecular descriptors and ECFPs (Extended Connectivity Fingerprints) from Dragon^17^ towards the development of a QSTR-based expert system^16,18^. BitterPredict^19^ enhanced the training data of BitterX^13^, by including molecules curated by Rojas et al.^18^ and those identified as being ‘probably non-bitter’ from Fenaroli’s Handbook of Flavor Ingredients^20^, while excluding the random molecules. They further established rigorous external validation sets (curated from literature and other unpublished resources), to show the efficacy of their AdaBoost model based on physicochemical and ADMET properties from Canvas. e-Bitter^21^ diverged from using random molecules as part of training data, and instead used only the experimentally verified bitter and non-bitter molecules along with ECFPs of varying lengths for training ensembles of machine learning models. BitterSweetForest^22^ is perhaps the only study till date to look at the dichotomy of bitter-sweet taste prediction, utilizing bitter, sweet compounds from BitterDB^14^ and SuperSweet^23^ respectively, towards the development of molecular fingerprints based Random Forest model.

While these studies have advanced our understanding of the molecular correlates of bitter-sweet taste and contributed predictive models, there is ample scope for improvement via a meticulous compilation of bitter-sweet molecules and utilization of a wide spectrum of molecular descriptors. The unavailability of trained models and datasets (in a timely fashion) as well as the use of proprietary software (such as Schrödinger, Dragon^17^) for generating molecular descriptors are major bottlenecks in making further advances. The only exception to this trend is e-Bitter^21^, which provides an end-to-end software, albeit only for bitter prediction. Further, these studies have relied on threshold-based metrics such as sensitivity and specificity-which are highly sensitive to the specific cut-offs used-to evaluate the performance of their models.

In this study, we aimed to create an integrative machine learning framework for bitter-sweet classification based on well-curated data, exhaustive set of molecular descriptors, and use of meaningful performance metrics. Towards this goal, we compiled an extensive dataset of structurally diverse bitter, non-bitter, sweet and non-sweet molecules, and used an array of 2D and 3D molecular descriptors compiled from both proprietary as well as open source software. Interestingly, models trained using open source ChemoPy descriptors exhibited performance at par with those trained using descriptors from proprietary software. Furthermore, all relevant features for bitter-sweet prediction were identified using the Boruta algorithm, which significantly reduced the dimensionality of the feature space. We present machine learning models implementing Random Forest, Ridge Logistic Regression and AdaBoost (decision trees) with performance matching or exceeding the state-of-the-art for, both, bitter and sweet classification. Importantly, we provide the datasets as well as the machine learning models with the hope that these will help in advancing the knowledge of the molecular basis of bitter-sweet taste and building relevant applications.

## 2. RESULTS

We addressed the problem of predicting bitter-sweet taste of a molecule by application of machine learning algorithms. For training and evaluating the models, we compiled an extensive set of molecules, and represented them via a range of molecular descriptors (open source as well as proprietary). Appropriate pre-processing techniques were applied to address redundancy in molecular descriptors. The performance of our models (BitterSweet) were evaluated using meaningful metrics and compared to the existing state-of-the-art models. Further, we also identified the specific features/feature blocks critical for bitter/non-bitter and sweet/non-sweet prediction, and enumerated the bitter-sweet chemical space of diverse sets of specialized compounds using BitterSweet models.

### 2.1 Data Compilation and Curation

Towards the creation of machine learning models for bitter-sweet prediction, previous studies have attempted compilation of data of molecules with bitter, sweet taste and appropriate controls (Table 1). However, inconsistencies in the curation process due to the inclusion of molecules with unverified taste information and incomplete representation of chemical space can lead to incorrect inferences and predictions. BitterPredict^19^ and BitterX^13^ have used unverified non-bitter molecules as a significant part of their training sets (55.6% and 50% respectively), which potentially adds noise to their models. While e-Bitter^21^, Rojas et al.^16^ and BitterSweetForest^22^ mitigated this problem by utilizing only experimentally verified data, this significantly reduced the size of their datasets and might have led to insufficient representation of the bitter-sweet chemical space. Hence we surmise that towards an effective model for prediction of bitter-sweet taste, an exhaustive compilation of bitter, non-bitter, sweet and non-sweet compounds to span the chemical space is essential while not compromising the accuracy of taste information of the molecules.

**Table 1:** Overview of previous research on bitter-sweet taste prediction with details of the type of molecular descriptor used, the source of data and control(s) used, the nature and size of these data.

| Reference | Molecular Descriptors | Taste | Source | Type | Number |
| --- | --- | --- | --- | --- | --- |
| BitterX<sup>13</sup> | Common Physicochemical Descriptors | Bitter | BitterDB<sup>14</sup> | Verified | 539 |
|  |  | Non-bitter | In-house data | Verified | 20 |
|  |  |  | Available Chemicals Directory | Random | 519 |
| Rojas et al.<sup>16</sup> | ECFPs and Dragon2D molecular descriptors | Sweet | Literature (Refer to supplementary materials of Rojas et al.<sup>16</sup>) | Verified | 336 |
|  |  | Bitter |  | Verified | 81 |
|  |  | Tasteless |  | Verified | 130 |
| BitterPredict<sup>19</sup> | Physicochemical and ADMET descriptors | Bitter | BitterDB<sup>14</sup> and Rojas et al.<sup>18</sup> (bitter) | Verified | 691 |
|  |  | Tasteless | Rojas et al.<sup>18</sup> (tasteless) | Verified | 130 |
|  |  | Probably non-bitter | Fenaroli's Handbook of Flavor Ingredients<sup>20</sup> | Random | 1451 |
|  |  | Sweet | Rojas et al.<sup>18</sup> (sweet) | Verified | 336 |
| e-Bitter<sup>21</sup> | ECFPs | Bitter | BitterDB<sup>14</sup>, Rodgers et al.<sup>12</sup> and Rojas et al.<sup>18</sup> (bitter) | Verified | 707 |
|  |  | Non-bitter | Rojas et al.<sup>18</sup> (sweet and tasteless) and BitterX<sup>13</sup> | Verified | 149 |
|  |  | Sweet | SuperSweet<sup>23</sup>, SweetenersDB<sup>43</sup> and Literature<sup>18,44,45</sup> | Verified | 443 |
| BitterSweetForest<sup>22</sup> | Morgan, Atom-Pair, Torsion and Morgan Feat fingerprints | Bitter | BitterDB<sup>14</sup> | Verified | 685 |
|  |  | Sweet | SuperSweet<sup>23</sup> | Verified | 517 |

In this study, we curated information on molecules with bitter-sweet taste from a wide variety of resources ranging from scientific publications to books. After removing molecules for which exact information of their bitter/sweet taste was either unavailable or conflicting, the curated dataset consisted of 918 bitter and 1510 non-bitter molecules as well as 1205 sweet and 1171 non-sweet molecules. Tasteless compounds were included as important controls for both bitter and sweet taste prediction. The datasets were split into training and testing sets such that the latter corresponded to the external validation/test sets established by BitterPredict^19^ for bitter/non-bitter prediction and Rojas et al.^16^ for sweet/non-sweet prediction (Supplementary Table S1). The curated training dataset is structurally diverse when seen in comparison to random bioactive molecules from ChEBI^24^, as evident in the 2D t-SNE plot generated using the physicochemical features (Figure 1), with molecules from different sources incrementally capturing subsets of the general chemical space.

**Figure 1:**
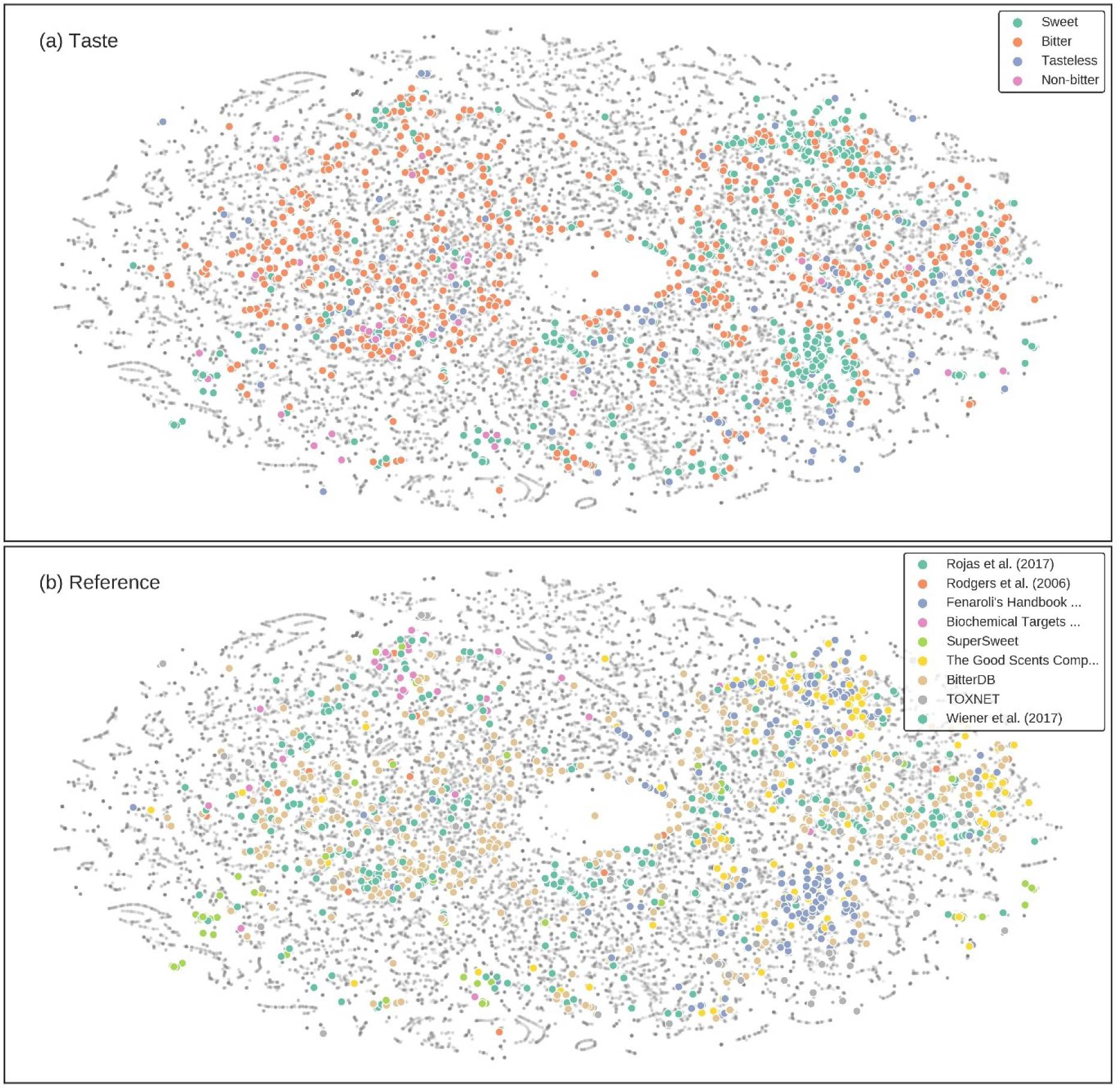
2D t-SNE scatterplot of curated and random molecules (ChEBI) generated using physiochemical features from Canvas. (a) Annotating taste information of molecules reveals the structural diversity of bitter, sweet and tasteless compounds as compared to random molecules. (b) Molecules from different sources incrementally capture subsets of the general chemical space.

### 2.2 Molecular Descriptors

Other than the quality of training and validation datasets, the choice of relevant features plays a key role in the performance of machine learning models. Over the years various molecular descriptors have been suggested to be relevant for bitter-sweet taste prediction (Table 1). While BitterX^13^ used physicochemical descriptors prescribed by the Handbook of Molecular Descriptors, BitterPredict^19^ used ADMET properties in addition to physicochemical features from Canvas. Rojas et. al^16^ relied on 2D molecular descriptors and ECFPs from Dragon. Both e-Bitter^21^ and BitterSweetForest^22^ used binary fingerprints, with the former using ECFPs and the latter resorting to an amalgamation of Morgan, Atom-Pair, Torsion, and Morgan Feat fingerprints. We intended to use an exhaustive set of descriptors from both commercial as well as open source software, to ascertain their contribution towards bitter-sweet taste of molecules. Towards this end, we implemented an array of features (2D/3D QSAR and ADMET descriptors, binary fingerprints) enumerating key aspects of molecular properties: Dragon 2D, Dragon2D/3D, Chemopy and Canvas. Supplementary Table S2 summarises the details of feature sets used in this study.

### 2.3 Feature Selection

The high-dimensionality and multicollinearity of certain molecular descriptor sets (Dragon2D, Dragon2D/3D, and ChemoPy) necessitated pre-processing to prune uninformative descriptors or to combine correlated features in a meaningful manner. We implemented the Boruta algorithm^25^ to find all the relevant features for bitter-sweet prediction. This led to significant reductions (35%-90%) in all the descriptor sets except Canvas (Figure 2). We also used Principal Component Analysis (PCA) to linearly combine attributes, such that the derived features captured the maximum variance in the data and were orthogonal to each other. Interestingly, more than 90% of the total variance could be captured by using the first three principal components in all of the descriptor sets (Supplementary Figure S1).

**Figure 2:**
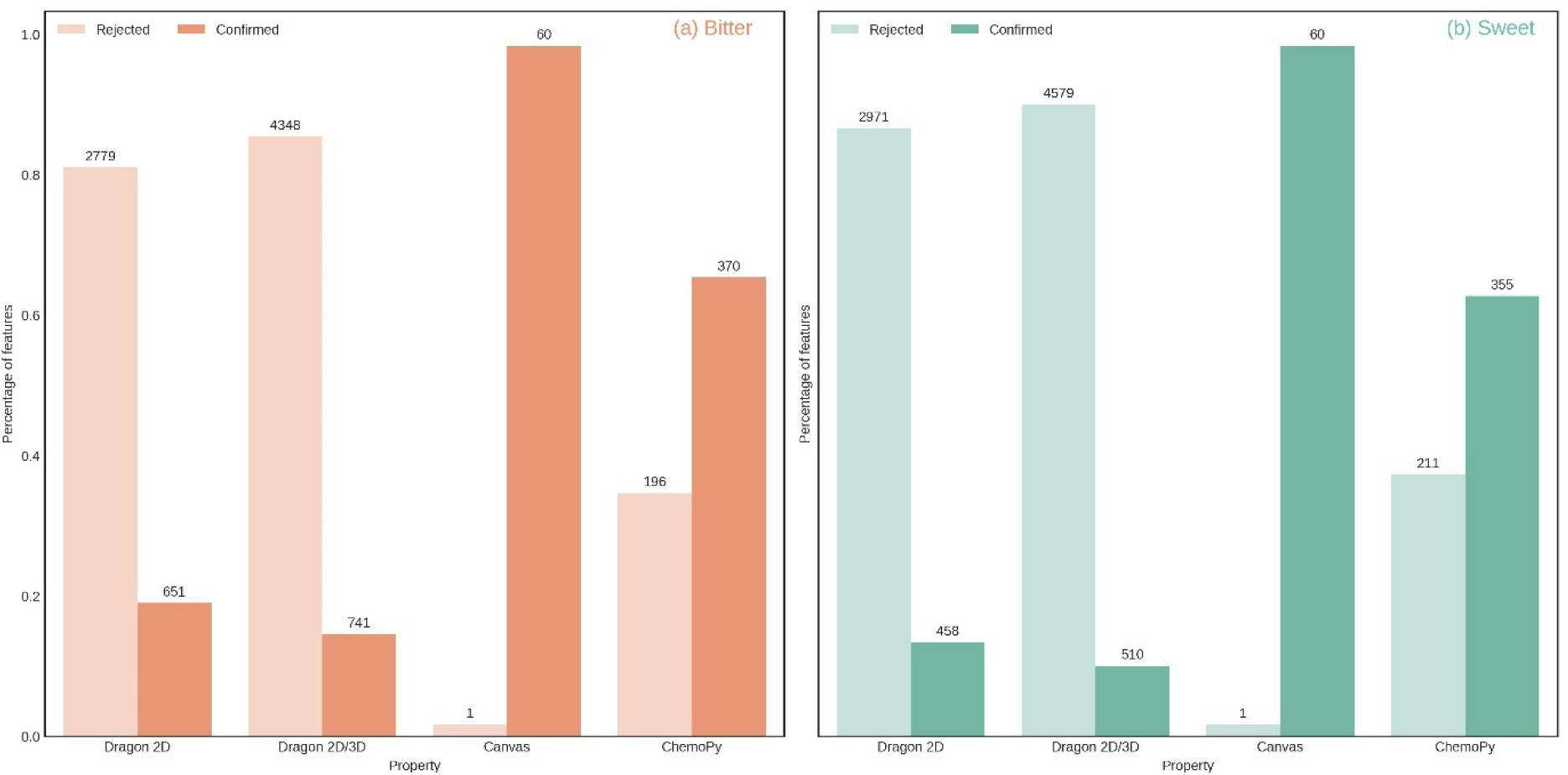
Percentage of features retained in Dragon2D, Dragon2D/3D, Canvas, and ChemoPy descriptor sets after the application of Boruta algorithm. For both (a) Bitter/Non-Bitter and (b) Sweet/Non-Sweet prediction datasets, a significant number of features from Dragon2D, Dragon2D/3D, and ChemoPy molecular descriptor sets were deemed as unimportant.

### 2.4 Model Performance and Comparison

After applying pre-processing on the curated data, Random Forest (RF), Ridge Logistic Regression (RLR) and Adaboost (AB) machine learning models were trained to classify a molecule as bitter/non-bitter and sweet/non-sweet, using each set of five molecular descriptors separately (Figure 3). The model parameters were calibrated using 5-fold stratified cross-validation. Henceforth, all our models are collectively referred to as ‘BitterSweet’. To avoid contingency of evaluation metrics on specific thresholds, BitterSweet models were evaluated using threshold-independent metrics such as Area Under Precision-Recall Curve (AUPR) and Area Under Receiver Operating Characteristic Curve (AUROC) in addition to F1-score, sensitivity, and specificity.

**Figure 3:**
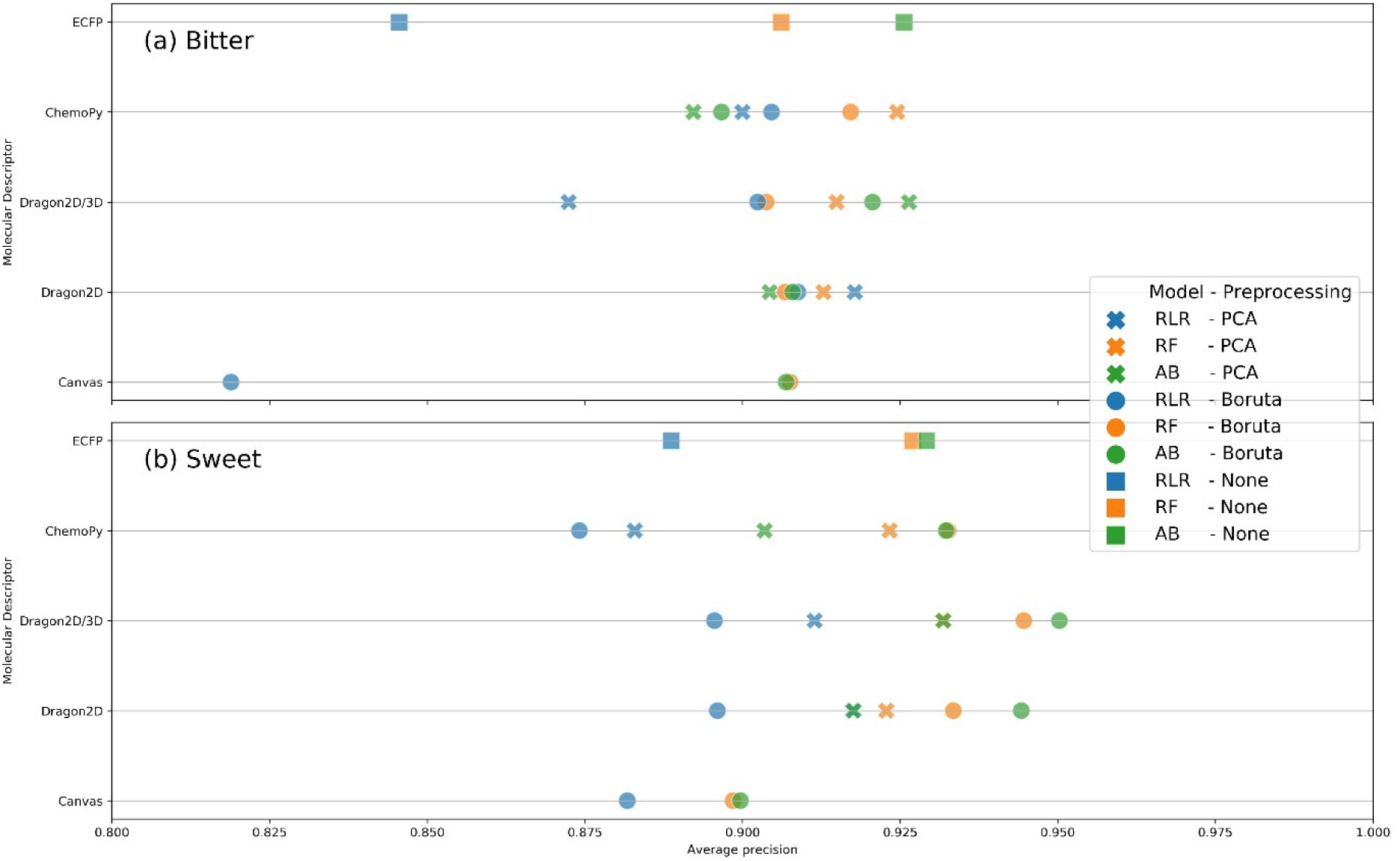
Performance (in terms of Average Precision) of the best BitterSweet models corresponding to each molecular descriptor set for (a) Sweet/Non-Sweet prediction and (b) Bitter/Non-Bitter prediction. Dragon2D/3D molecular descriptor set and Boruta feature selection were found to produce the most optimal models for sweet/non-sweet prediction. For bitter/non-bitter prediction, PCA outperformed Boruta.

**Sweet/Non-Sweet Prediction**: Fig 3(b) depicts performance (AUPR score) of BitterSweet models utilizing different molecular descriptor sets, algorithms, and pre-processing methods. Adaboost and Random Forest models trained after application of Boruta algorithm were found to outperform other models consistently. PCA performed better than Boruta only in the case of Ridge Logistic Regression. While Dragon2D/3D molecular descriptor set was found to give the best performance, Dragon2D and ChemoPy were marginally worse. On the other hand, models trained using Canvas descriptors performed the worst by a significant margin, despite 3D conformers being used for computation. Remarkably, models trained using (relatively simpler molecular descriptors) ECFPs performed were found to be better than those based on Canvas, and almost equivalent in performance to the ones using ChemoPy descriptors. A detailed performance profile of BitterSweet models on Cross-Validation and Test datasets in terms of each of the metrics (AUPR, AUROC, F1, NER, Sensitivity and Specificity) is provided in Supplementary Table S3.

**Bitter/Non-Bitter Prediction**: Fig 3(a) elucidates the performance (AUPR score) of BitterSweet models utilizing different molecular descriptor sets, algorithms, and pre-processing methods. There were a few differences compared to sweet/non-sweet classification. Ridge Logistic Regression was no longer the worst model, even achieving the best and second-best score in ChemoPy and Dragon2D molecular descriptors set. Random Forest was found to be competitive across all datasets. Regarding pre-processing methods, PCA consistently outperformed Boruta in contrast to sweet/non-sweet prediction, where the converse was true. Remarkably, models trained using the open source ChemoPy descriptors achieved the best performance. However, their improvement over ECFPs, Dragon2D, and Dragon2D/3D molecular descriptor based models was marginal. Canvas descriptors resulted in the worst models, despite using 3D conformers for computation. Detailed performance profiles of BitterSweet models in terms of each of the metrics (AUPR, AUROC, F1, NER, Sensitivity and Specificity) on Cross-Validation and Test datasets are provided in Supplementary Table S4, S5, S6 and S7 respectively.

**Comparison with Previous Studies**: While we emphasize on the need to use threshold-independent metrics in order to mitigate contingency on specific thresholds, a lack of vigilance in this regard by previous studies compels us to provide comparisons using non-error rate (NER), sensitivity (Sn), specificity(Sp) and F1-score. Table 2 and Table 3 list the performance of the best model corresponding to each molecular descriptor set for sweet/non-sweet and bitter/non-bitter prediction respectively, and their comparison with previous studies.

**Table 2:** Comparison of performance of the best BitterSweet models (corresponding to each molecular descriptor set) with Rojas et al.^16^ on the sweet/non-sweet test set. NA (Non-Availability) refers to the percentage of molecules in the test for which no predictions were made. The QSTR-based expert system of Rojas et al.^16^ combined structural similarity analysis based on ECFPs with N-nearest neighbors (N3) and partial least squares discriminant analysis (PLSDA classifiers).

|  | Molecular Descriptors | Model | AUPR | AUROC | F1 | NER | Sn | Sp | NA |
| --- | --- | --- | --- | --- | --- | --- | --- | --- | --- |
| BitterSweet | Canvas | AB-Boruta | 0.900 | 0.837 | 0.791 | 0.7710 | 0.705 | 0.837 | 4.4% |
|  | Dragon2D | AB-Boruta | 0.944 | 0.881 | 0.829 | 0.7995 | 0.769 | 0.830 | 2.5% |
|  | Dragon2D/3D | AB-Boruta | 0.950 | 0.883 | 0.856 | 0.8340 | 0.790 | 0.878 | 4.4% |
|  | ChemoPy | RF-Boruta | 0.933 | 0.852 | 0.772 | 0.8005 | 0.641 | 0.960 | 5% |
|  | ECFPs | AB | 0.929 | 0.847 | 0.806 | 0.7875 | 0.726 | 0.849 | 1.24% |
| Rojas et al.<sup>16</sup> | ECFPs & Dragon2D | QSTR-based expert system | - | - | - | 0.848 | 0.880 | 0.816 | 19.3% |

**Table 3:** Comparison of performance on the bitter/non-bitter test sets of the best BitterSweet models (corresponding to each molecular descriptor set) with BitterX, BitterPredict, and e-Bitter.

|  | Molecular Descriptors | Model | Phyto-Dictionary |  |  |  | UNIMI |  |  |  | Bitter-New |  |  |  |
| --- | --- | --- | --- | --- | --- | --- | --- | --- | --- | --- | --- | --- | --- | --- |
|  |  |  | Sn | Sp | F1 | AUPR | Sn | Sp | F1 | AUPR | Sn | Sp | F1 | AUPR |
| BitterSweet | Canvas | RF-Boruta | 0.980 | 0.76 | 0.932 | 0.959 | 0.826 | 0.727 | 0.745 | 0.810 | 0.652 | - | 0.789 | - |
|  | Dragon2D | RLR-PCA | 0.979 | 0.64 | 0.904 | 0.970 | 0.609 | 0.970 | 0.737 | 0.847 | 0.609 | - | 0.757 | - |
|  | Dragon2D/3D | RF-PCA | 0.980 | 0.68 | 0.914 | 0.977 | 0.913 | 0.636 | 0.750 | 0.746 | 0.913 | - | 0.955 | - |
|  | ChemoPy | RLR-PCA | 0.939 | 0.68 | 0.893 | 0.957 | 0.870 | 0.818 | 0.816 | 0.864 | 0.696 | - | 0.821 | - |
|  | ECFPs | AB | 0.959 | 0.84 | 0.940 | 0.970 | 0.913 | 0.697 | 0.778 | 0.783 | 0.652 | - | 0.789 | - |
| BitterX<sup>13</sup> | Handbook of Molecular Descriptors | SVM | 0.939 | 0.308 | 0.814 | - | 0.652 | 0.562 | 0.577 | - | 0.739 | - | 0.850 | - |
| BitterPredict<sup>19</sup> | Canvas | AB | 0.980 | 0.692 | 0.914 | - | 0.783 | 0.848 | 0.783 | - | 0.739 | - | 0.850 | - |
| e-Bitter<sup>21</sup> | ECFPs | CM01 | 0.980 | 0.769 | 0.932 | - | 0.913 | 0.545 | 0.712 | - | 1.000 | - | 1.000 | - |

For the sweet-prediction task, BitterSweet models were only compared with the QSTR-based expert system developed in Rojas et al.^16^ due to the unavailability of BitterSweetForest^22^ model as well as differences in external test sets (Table 2). While their QSTR-system achieved an impressive NER of 0.848, it was unable to make predictions for 30 (19.3%) molecules in the test set. On the contrary, BitterSweet models were able to predict the sweet taste (along with probability values) of all molecules in the test except for a small fraction for which molecular descriptors could not be generated (due to incomplete structures). Dragon2D/3D descriptors based AB-Boruta model achieved the best NER score of 0.834 while leaving out only 7 (4.4%) test set molecules. Similarly, AB-Boruta models based on ECFPs, Dragon2D, ChemoPy, and Canvas descriptors achieved competitive NER scores of 0.788, 0.8, 0.801, and 0.791 respectively while ignoring just 2-7 molecules.

For the task of bitter-prediction, we provide performance comparison with BitterX, BitterPredict, and e-Bitter (Table 3). On the Phyto-dictionary test set, AB model based on ECFPs was presented with the best F1-score (0.94) and outperformed e-Bitter, BitterPredict and BitterX by a margin of 0.8%, 2.6%, and 11.6% respectively. Further, it attained an AUPR value of 0.97, suggesting that the classifier has almost perfect performance for Phyto-dictionary molecule set. Other BitterSweet models were also found to fare well in comparison, achieving F1 scores in the range 0.89-0.94 and AUPR scores around 0.96-0.98. UNIMI set was found to be more challenging than other validation sets due to the presence of molecules with similar scaffolds but different tastes. While the RLR-PCA model based on ChemoPy descriptors achieved the best F1-score (0.816) and exceeded the F1-scores achieved by e-Bitter (0.712), BitterPredict(0.783), and BitterX (0.577 F1-score), other BitterSweet models were also found to be competitive (0.737-0.778). The Bitter-new molecule set was the smallest bitter/non-bitter test set, comprising of just 23 bitter molecules. Lack of presence of non-bitter molecules made the calculation of all metrics besides sensitivity infeasible. In addition, the relatively small size of this set resulted in an overall 4.4% increase/decrease based on the outcome of prediction of a single molecule. Among our models, RF-PCA attained the best sensitivity of 0.913 exceeding both BitterPredict and BitterX, while falling just a little short of the perfect sensitivity achieved by e-Bitter. However, other BitterSweet models were significantly inferior with sensitivity scores in the range 0.609-0.696.

In summary, the best BitterSweet models had a significantly larger applicability domain as compared to Rojas et al.’s QSTR-based expert system^16^ while achieving similar performance. They were also found to be robust to class imbalance–in contrast to BitterX^13^ and e-Bitter^21^– both of which had a larger number of false positives for Phyto-dictionary and UNIMI validation sets respectively. In addition, BitterSweet models are applicable for prediction of both bitter and sweet tastes, and provide probabilities (as opposed to only dichotomous classification) as a measure of confidence.

### 2.5 Feature Importance

Calculation of molecular descriptors is a computationally intensive process (especially 3D properties), with inadequacies in the specification of compounds’ structures being an additional hindrance. Information regarding relevant features and feature types (blocks) can help in the better use of computational resources and expedite model development. One reliable measure of feature importance is the mean decrease in impurity (Gini impurity), obtained when using tree-based classifiers such as Random Forest. However, these importance values can be misleading in the presence of multicollinearity, where the true importance scores get shared between correlated features. Moreover, as opposed to the general objective of finding the shortest subset of most informative features in machine learning methods, constraints on the availability of molecular descriptors necessitates identification of ‘all’ relevant features. Towards these goals, the Boruta ‘all relevant feature selection’ algorithm (which is robust to the presence of multicollinearity) was used for identification of distinguishing features in our studies.

In order to find the relevance of feature blocks, the importance scores of their constituent features were aggregated (Figure 4). Among ChemoPy features, the descriptors associated with Charge and Bcut feature blocks were dominant for both, bitter/non-bitter and sweet/non-sweet prediction. As also seen at the level of individual ChemoPy descriptors (Supplementary Figure S2), a significant number of features belong to these blocks; ten out of the thirty most important features for bitter prediction and sixteen out of the thirty most important features for sweet prediction. Among the Dragon based molecular descriptors CATS2D, CATS3D, Molecular, and Constitutional feature blocks were among the most dominant blocks towards prediction of bitter-sweet taste (Figure 5, Supplementary Figure S3). In case of the molecular descriptors obtained through the Canvas software, in the absence feature blocks, we identified individual feature scores highlighting the importance of QPlogBB, FOSA, and QPlogbw among others (Supplementary Figure S4).

**Figure 4:**
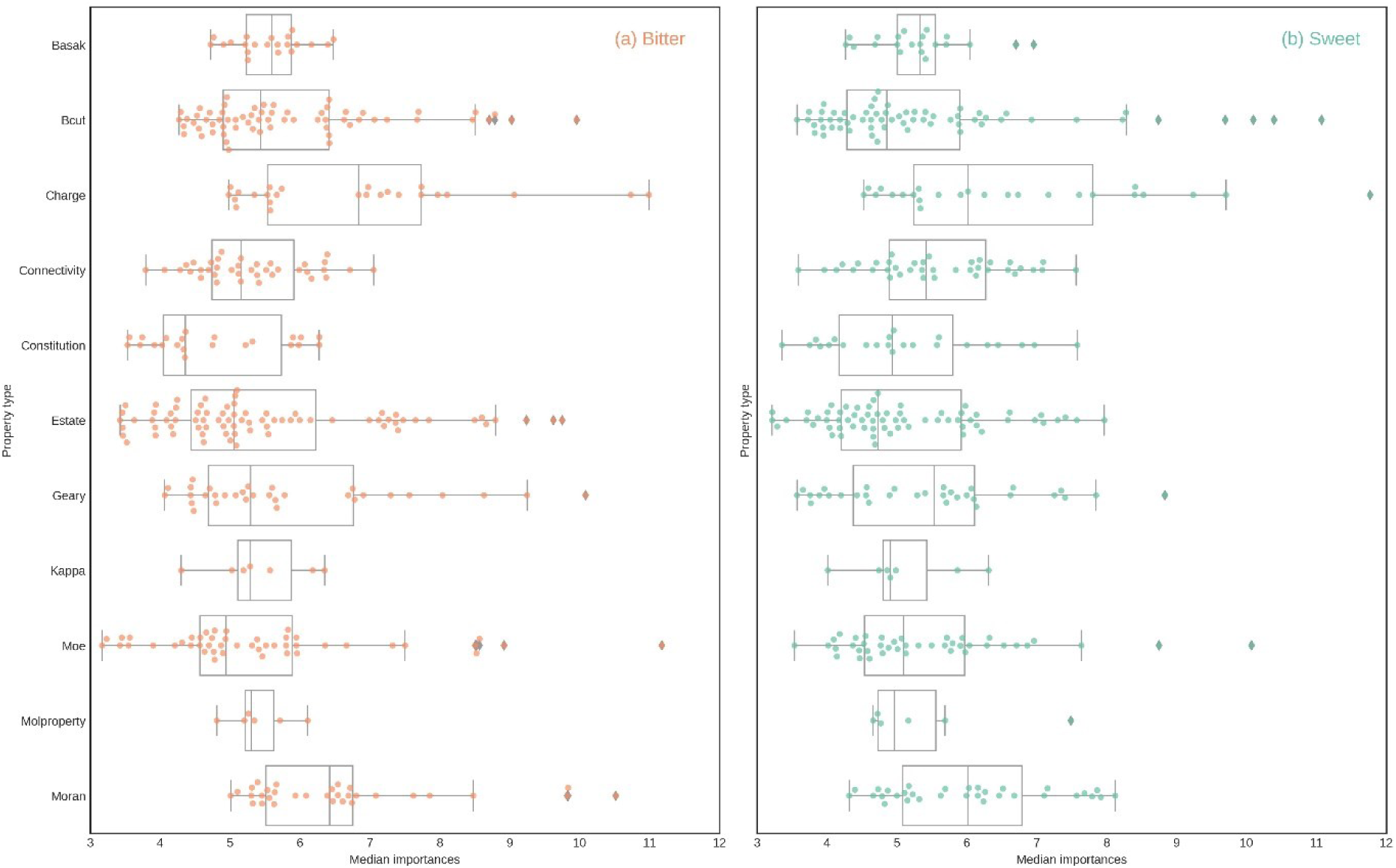
Boxplot of feature importance for each property block in ChemoPy set.

**Figure 5:**
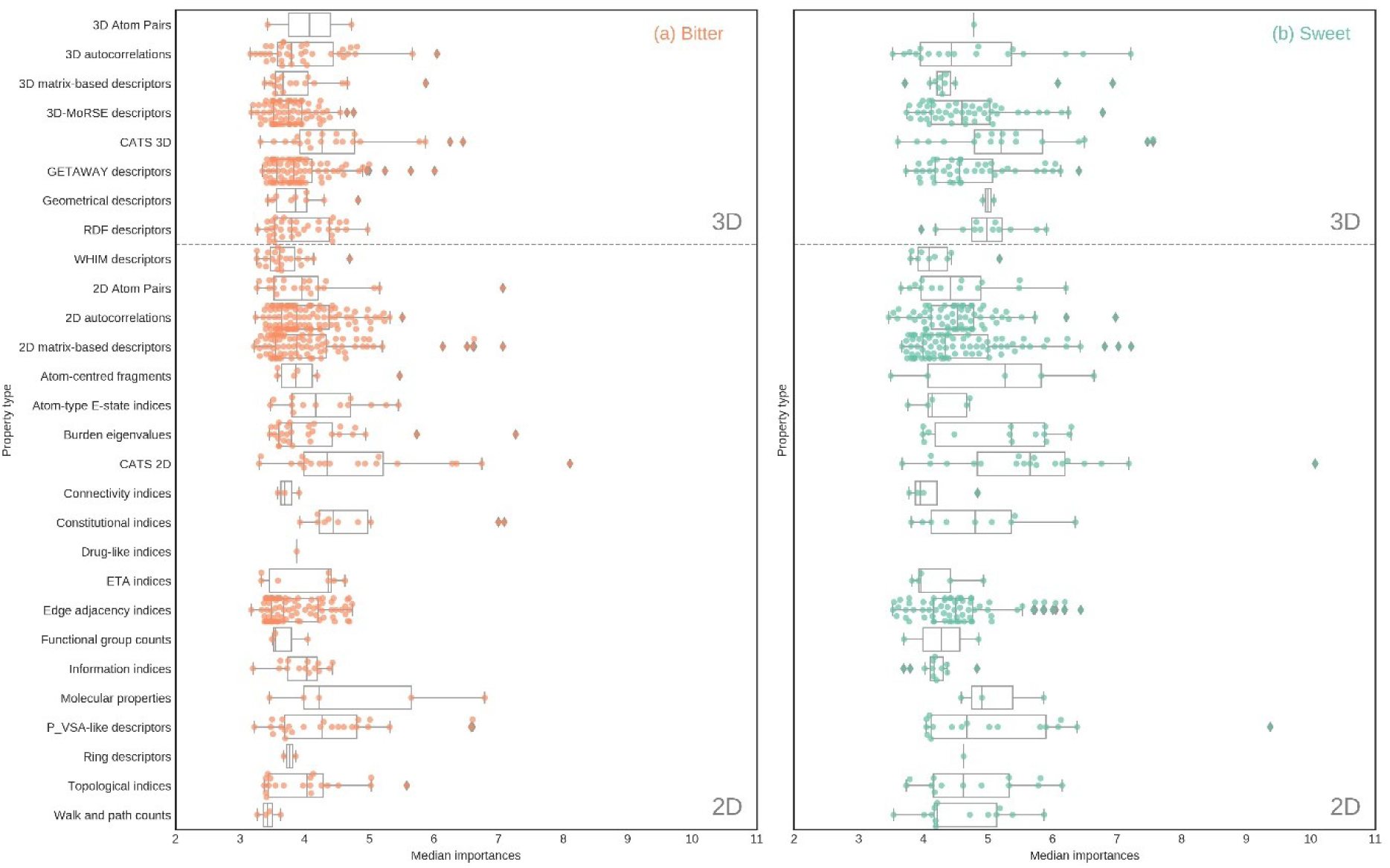
Boxplot of feature importance for each property block in Dragon 2D/3D set.

### 2.6 Applicability Domain Assessment

While making predictions, it is important to identify and prune molecules significantly different from the ones used as part of the training set in order to ensure consistent predictive performance. The Applicability Domain for BitterSweet models was defined in accordance with the guidelines set by the Organization of Economic Cooperation and Development. An unseen molecule is categorized as falling outside the applicability domain of a classifier, if its median Euclidean distance from the *N* most similar compounds in training set exceeds a threshold *δ*. The distance is found using only the *k* most important features as defined by the classifier. For the open source ChemoPy descriptors based Boruta models, we set the values for *N, δ* and *k* to be 5, 3 and 25. These parameter values were determined empirically on the basis of pairwise similarity of molecules in the training set and are not meant to be rigid. To achieve higher confidence in model predictions stricter thresholds may be used.

### 2.7 Model Application

In order to explore the dichotomy of bitter and sweet tastes of molecules in specialized chemical sets, the ChemoPy-based RF-PCA BitterSweet models were applied to databases of sweet (SuperSweet^23^), flavor (FlavorDB^26^), food (FooDB; http://foodb.ca), toxic (DSSTox^27^), natural (Super Natural II^28^), and drug (DrugBank^29^) molecules. The cutoffs for categorizing a molecule as bitter-sweet were set with the objective of balancing sensitivity and specificity on the cross-validation sets. Supplementary Figure S5 demonstrates the number of predicted bitter and sweet compounds for the above-mentioned databases at different cutoffs.

**SuperSweet**^23^: It comprises of more than 20000 artificial and natural molecules, among which a significant proportion are categorized as sweet-like, i.e., speculated ‘sweet’ molecules identified based on their structural similarity with 200 verified sweet compounds. Prediction for bitter-sweet taste was made for 18122 sweet-like molecules within the applicability domain of the models. Despite the high structural similarity of sweet-like compounds to sweet compounds, 4303 compounds (24%) were predicted to be bitter in contrast to 13639 (75%) sweet predictions. This is indicative of the fact that small changes in structure can radically alter taste perception.

**FlavorDB**^26^: The bitter-sweet predictions were made for the 2294 flavor molecules linked to natural sources in FlavorDB. Molecules outside the applicability domain of the models were removed, resulting in a reduced set of 2103 compounds. Among these, our models predicted 78% (1650 compounds) to be sweet, and 20% (416 compounds) to be bitter. This is interesting since most of these molecules are small and volatile odorous molecules modulated by approximately 400 olfactory receptors, some of which (class A family of GPCRs) share structural features with bitter receptors (binding pocket of TMD domain of *h* Tas1R).

**Super Natural II**^28^: Application of BitterSweet models on the 325282 natural molecules in Super Natural II can help enumerate the chemical space of natural bitter-sweet compounds. Among the 280989 molecules within the applicability domain of the models, 21% (59414) were predicted to be sweet whereas 62% (173215) were predicted to be bitter. This indicates that a significant proportion of natural molecules taste bitter.

**FooDB**: It contains 26319 food molecules, among which 20122 were within the applicability domain of the models. Similar to the BitterPredict model, a significant number of molecules in FooDB (38%; 7560) were predicted to be bitter possibly due to the presence of bitter-tasting glucosinolates, terpenes, flavonoids and alkaloids in plants. In contrast, 42% of compounds (8525) were predicted to be sweet, suggesting that the chemical space of food compounds has bitter and sweet molecules in almost equal proportions.

**DSSTox**^27^: It is widely assumed that bitter taste modality was evolved to preclude the consumption of toxic compounds^30-32^, with a large number of toxic compounds known taste bitter. The Distributed Structure-Searchable Toxicity Database (DSSTox) consists of 719795 toxic compounds, among which 580606 compounds were within the applicability domain of the models. 55% (319463) of these compounds were predicted to be bitter in contrast to 26% (148187) sweet predictions consistent with the assumption. However, as evident from the large number of sweet and non-bitter predictions, bitterness cannot be considered a reliable beacon for toxicity.

**DrugBank**^29^: The bitter-sweet taste prediction was made for 7049 molecules (among 8827) categorized as ‘approved small molecule drug’ and/or ‘experimental drug’ present within the applicability domain of our models. Similar to BitterPredict model, a significant number of molecules were predicted to be bitter (63%; 4426) in contrast to 22% (1493) sweet predictions. This result is consistent with the evidence prevalence of bitter taste among drugs^33,34^.

### 2.8 Software for BitterSweet prediction

We provide all BitterSweet models, datasets used for training them as well as an end-to-end pipeline (https://github.com/cosylabiiit/bittersweet/) for prediction of bitter-sweet taste of molecules. The pipeline relies on freely available ChemoPy molecular descriptors and the state-of-the-art RF-PCA models. The software is freely available for non-commercial use. A BitterSweet prediction server along with a user-friendly interface for exploring these data are also being made available^35^.

## 3. Discussion

In this study, we have established the largest dataset of verified bitter, sweet and tasteless compounds till date. Using these data, we have identified the most relevant features and feature blocks in diverse molecular descriptor sets (2D, 3D, open source and proprietary) and trained bitter-sweet prediction models with performance comparable to (and in some cases exceeding) the state-of-the-art. Given the complexity of taste prediction, while one may expect more nuanced descriptors to play a critical role in taste prediction, we observed that the performance of models trained using 2D descriptors was not significantly different from those trained using 3D descriptors. Remarkably, BitterSweet models trained using open source ChemoPy descriptors achieve performance comparable to those trained using molecular descriptors from proprietary software for both bitter/non-bitter and sweet/non-sweet prediction. Finally, we would like to highlight the importance of using threshold-independent performance metrics for meaningful assessment of models. We hope that future studies would consistently use these metrics so as to enable better evaluation of their models. Application of BitterSweet models on large specialized chemical sets suggested that a significant proportion of natural, toxic and drug-like molecules are bitter. On the contrary, natural flavor molecules were largely predicted to be sweet while bitter-sweet prediction for food molecules suggested the presence of an equal number of bitter and sweet compounds. BitterSweet models can be of immense value towards the objective of selection and synthesis of compounds with the desired gradient of bitter-sweet taste. And with that in mind, we release a state-of-the-art bitter-sweet prediction tool based on freely available ChemoPy descriptors. Despite curating the largest dataset of bitter/non-bitter and sweet/non-sweet molecules as well as using diverse molecular descriptors set, our models do not dramatically improve upon the existing ones. However, this might be a limitation of the structural diversity of bitter-sweet molecules represented by the test set, and future studies should evaluate their relevance as a gold standard.

Taste is a complex, multifactorial sensation. Lack of datasets concerning the intensity of bitterness/sweetness and other primary tastes (salty, sour and umami) of molecules hinders the development of nuanced taste prediction models. While BitterSweet models can be utilized for individual compounds, predicting the taste of a mixture of compounds (natural or artificial) may be of more practical value. However, at present the utility of BitterSweet models is limited, and they cannot be directly used for aggregate taste prediction of a mixture of compounds, where concentration and competitive binding to taste receptors are key factors. Despite an extensive compilation of bitter and sweet molecules along with appropriate controls for training BitterSweet models, the reliability of predictions for unseen molecules is constrained by the structural diversity of the training set (See ‘Applicability Domain Assessment’ section). We hope that with access to nuanced data, future studies will address some of these challenges.

To the best of our knowledge, this is the first study which compares and contrasts the performance of diverse molecular descriptor sets for bitter-sweet prediction and releases pre-trained models. We believe that along with computational strategies for generating realizable small organic molecules^36^, our models would provide the foundation of a framework to span the chemical universe in search of compounds with desirable bitter-sweet taste gradient.

## 4. MATERIALS & METHODS

### 4.1 Data compilation and curation

The data of bitter-sweet molecules was curated from various pre-existing databases, books, and research articles. Incorporation of structurally diverse molecules from different sources ensured a meaningful representation of the chemical space for training as well as evaluating the machine learning models. A brief summary of the datasets is given in Table 4. The following resources were used: (a) Biochemical Targets of Plant Bioactive Compounds by Gideon Polya^15^; (b) BitterDB^14^; (c) Fenaroli’s Handbook of Flavor Ingredients^20^ (5th Edition); (d) Rodgers et al. (2006)^12^; (e) Rojas et al. (2017)^16^; (f) SuperSweet^23^; (g) TOXNET^37^; (h) The Good Scents Company Database (www.thegoodscentscompany.com); and (i) BitterPredict^19^. The chemical structures of molecules belonging to (c), (g) and (h) were obtained via a two-step process. First, their Chemical Abstract Services numbers were converted to IUPAC International Chemical Identifier (InChI) using NIH’s Chemical Identifier Resolver (https://cactus.nci.nih.gov/chemical/structure). The InChIs were then used to retrieve chemical structures from PubChem. For obtaining chemical structures of molecules from (a), manual lookup was performed on PubChem. The general set of biologically reactive molecules from ChEBI was designated as the random set and used for performing comparative experiments with bitter-sweet molecules.

**Table 4:** Overview of the resources used for creating bitter/non-bitter and sweet/non-sweet datasets.

| S. No. | Reference | Taste | Number of curated molecules |
| --- | --- | --- | --- |
| (a) | Biochemical Targets of Plant Bioactive Compounds by Gideon Polya<sup>15</sup> | Bitter | 39 |
|  |  | Sweet | 32 |
| (b) | BitterDB<sup>14</sup> | Bitter | 592 |
| (c) | Fenaroli's Handbook of Flavor Ingredients (5th Edition)<sup>20</sup> | Bitter | 33 |
|  |  | Sweet | 426 |
|  |  | Tasteless | 3 |
| (d) | Rodgers et al. (2006)<sup>12</sup> | Bitter | 29 |
| (e) | Rojas et al. (2017)<sup>16</sup> | Bitter | 81 |
|  |  | Sweet | 433 |
|  |  | Tasteless | 133 |
| (f) | SuperSweet<sup>23</sup> | Sweet | 198 |
| (g) | TOXNET<sup>37</sup> | Tasteless | 72 |
| (h) | The Good Scents Company Database (www.thegoodscentscompany.com) | Bitter | 43 |
|  |  | Sweet | 158 |
| (i) | Wiener et al. (2017)<sup>19</sup> | Bitter | 105 |
|  |  | Non-bitter | 66 |

**Bitter/Non-Bitter Data**: The references numbered (a)-(h) in Table 4 were used to create the training set of the bitter-taste prediction model. The positive set comprised of ‘bitter’ molecules (813) whereas the negative set consisted of ‘sweet’ or ‘tasteless’ molecules (1444). The data curated from BitterPredict^19^ (i), comprising of (105) ‘bitter’ and (66) ‘non-bitter’ molecules was designated as the test set (Phyto-dictionary, UNIMI, Bitter-new), and used to facilitate comparison with previous studies (Supplementary Table S1).

**Sweet/Non-Sweet Data**: The training set for the sweet-taste prediction model consisted of molecules from (a)-(i) mentioned in Table 4, with only the ‘training’ subset being used from (e). While the negative set consisted of ‘bitter’ and ‘tasteless’ molecules (1066), the positive set comprised of ‘sweet’ molecules (1139). The ‘testing’ subset from (e) was designated as the test set to facilitate comparison with previous studies (Supplementary Table S1).

### 4.2 Processing the molecules

Canonical SMILES were obtained for all molecules using OpenBabel^38^, and subsequently used to prune duplicate structures. To further reduce noise, peptides, salt ions and molecules with less than 3 atoms were removed. Epik^39^ and LigPrep [Schrödinger Release 2018-3: LigPrep, Schrödinger, LLC, New York, NY, 2018] were used to obtain 3D conformers and protonation states of molecules at biological pH 7±0.5. If specified, the original chirality of the molecule was maintained. Finally, only the conformer with the lowest energy was retained for each molecule.

### 4.3 Molecular Descriptors

The bitter-sweet taste prediction models were trained and evaluated using Dragon 2D/3D QSAR descriptors^17^ and Extended Connectivity Fingerprints (ECFPs), physicochemical as well as ADMET (Absorption, Distribution, Metabolism, Excretion, and Toxicity) properties from Canvas^40^, and structural and physicochemical descriptors from ChemoPy^41^.

**Dragon 2D/3D QSAR Descriptors:** Central to the efficacy of Dragon software is its unique algorithm, which enables calculation of descriptors even in the presence of disconnected structures (by appropriately modifying the computational procedure). The SMILES string of molecules was used to compute the 2D molecular descriptor set, whereas 3D conformers and tautomers were utilized towards the calculation of 2D and 3D molecules descriptor set. The Dragon 7 software is commercially available.

**ECFPs (Extended Connectivity Fingerprints)**: ECFPs are circular fingerprints developed for structure-activity modeling, via which each molecule is represented as a binary vector with a predetermined number of bits and a maximum pattern length. Each feature in the binary vector denotes presence or absence of a particular substructure. Starting with molecular SMILES, 2048 bits ECFPs (2 bits per structural patterns and a maximum pattern length of 2) were computed using the commercially available Dragon 7 software.

**Canvas Descriptors**: Using the 3D conformers and tautomers of molecules, certain Physicochemical and ADMET properties were calculated using commercially available Canvas software.

**ChemoPy**: Common structural and physicochemical descriptors are implemented in the open source Python-based ChemoPy software. These were computed using the SMILES string of molecules.

### 4.4 Evaluation Metrics and Models

A binary classifier yields four primary measures: True Positives (TP) – Number of positive instances correctly predicted; False Positives (FP) – Number of negative instances incorrectly predicted as positive; True Negatives (TN) – Number of negative instances correctly predicted; and False Negatives (FN) – Number of positive instances incorrectly predicted as negative. The following metrics were used to assess the performance of BitterSweet models:

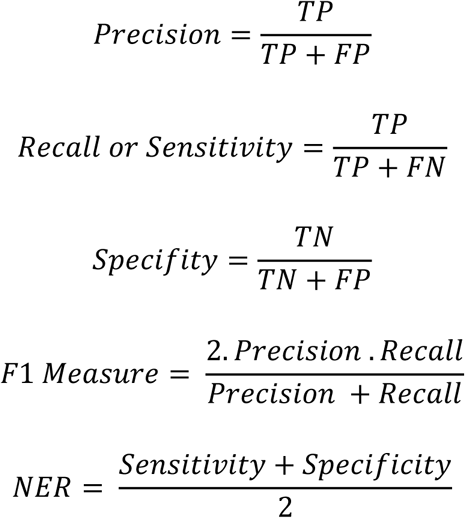

Area Under Curve – Receiver Operating Characteristic, is computed by evaluating *recall* and *fall-out (1 – specificity)* on a range of different threshold values.

**Ridge Logistic Regression**: Unlike standard regression, logistic regression tries to predict the ‘probability’ of a given input belonging to a particular class (bitter or non-bitter, sweet or non-sweet). The output always lies between [0,1]. Logistic regression involves a linear discriminant (separate data through linear boundary) and hence, serves as a good baseline. Its working involves maximizing a log-likelihood function or minimizing negative log likelihood (NLL). In the case of ‘Ridge’ logistic regression, in addition to minimizing NLL, a penalty is also imposed on the loss for regularization.

**Random Forest (RF)**: Random Forest algorithm is a type of ensemble learning method that utilizes bagged decision trees. They are quite versatile and can be used for both classification as well as regression. RF works by building a number of decision trees (usually greater than 100) at training time, each utilizing a subset of features and data points. At the time of prediction, the predictions made by its constituent decision trees are aggregated.

**Adaptive Boosting (AB)**: Boosting methods are a powerful way to enhance classifier performance and give state-of-the-art results on a variety of datasets. As opposed to ‘bagging’ where both the data points and features are sampled from the original data, boosting involves the sequential production of multiple learners, each attempting to correct errors from the previous learner. In this study we used Adaptive Boosting algorithm with decision trees as base learner.

### 4.5 Dimensionality Reduction

T-distributed Stochastic Neighbour Embedding (t-SNE): t-SNE^42^ is non-linear dimensionality reduction technique, particularly well-suited for visualizing high-dimensional datasets. The core idea behind t-SNE is to embed high-dimensional data into lower-dimensions in such a manner that the distance between dissimilar points is maximized and those between similar objects is minimized.

Principal Component Analysis (PCA): PCA is a mathematical technique that captures the linear interactions between the underlying attributes in the dataset. Every principal component can be expressed as a combination of one or more existing variables. All principal components are orthogonal to each other, and each one captures some amount of variance in the data.

### 4.6 Features Importance

Random Forest Relative Feature Importance: Every node in the ensemble of decision trees generated by the random forest algorithm is associated with a purity metric (Gini impurity). As the tree grows, this impurity value decreases. Nodes with the greatest decrease in the impurity metric occur at the start of the trees, while the ones with the least decrease occur at the end. By ranking features based on this impurity values, we are able to generate relative feature importance of the attributes used to make the prediction.

Selection of All Relevant Features Using Boruta Algorithm: The Boruta algorithm works on top of Random Forest classification algorithm. It captures the basic idea of ‘impurity metric’ in addition to the following steps to capture the feature importance: Duplicate the dataset and shuffle values in each column and call these values as shadow features; Train a standard Random Forest classifier over this new dataset; Check whether the ‘real’ features have a higher feature importance than ‘shadow’ features by evaluating the Z-Score metric; and at every iteration, compare the Z-scores and mark a feature as important if it is better than its shadow copies.

## Acknowledgements

The authors thank Indraprastha Institute of Information Technology (IIIT-Delhi) for providing computational facilities and support.

## Author Contributions Statement

G. B. and R.T. designed the study. R.T. curated the data. S.W., R.T. performed feature selection and importance ranking experiments, and trained the models. R.T. generated the bitter-sweet predictions for specialized chemicals sets. All the authors analysed the results and wrote the manuscript.

## Competing Interests

The authors declare that they have no competing interests.

## Data Availability

The datasets used for training and evaluating the BitterSweet models are available at https://github.com/cosylabiiit/bittersweet/data/.

